# Magnetoencephalography reveals adaptive neural reorganization maintaining lexical-semantic proficiency in healthy aging

**DOI:** 10.64898/2025.12.19.695466

**Authors:** Pietari Nurmi, Heidi Ala-Salomäki, Hanna Renvall, Mia Liljeström

## Abstract

Although semantic cognition remains behaviorally stable with age, neuroimaging studies report age-related alterations in response to semantic context. We aimed to reconcile these inconsistent findings by combining behavioral and magnetoencephalography (MEG) measures and examined the neural mechanisms underlying lexical-semantic processing in healthy aging. Forty-six Finnish-speaking adults (25 female) across younger (22– 32 years) and older (62 – 69 years) cohorts performed a semantic congruency judgment task on word pairs. Both age groups performed the task with high accuracy, with older adults responding faster than young. While advancing age was not associated with impaired task performance, MEG revealed neural reorganization in the old compared to young adults. Young participants exhibited the canonical N400 effect in the left temporal cortex 300 – 400 ms post-stimulus, followed by a Late Positive Complex (LPC), whereas older adults showed less pronounced congruency effects, and the activation distribution was shifted toward frontal and bilateral regions. Despite the absence of the canonical N400 effect in older participants, multivariate time-resolved decoding demonstrated that the congruency effect was decodable from the MEG responses in both groups from ∼300 ms post-stimulus, indicating preserved sensitivity to congruency with age. Decoding performance correlated with response time, suggesting that increased neural separation between semantic concepts facilitates rapid decision-making. Critically, age groups were distinguishable based on their neural responses 300 – 800 ms post-stimulus, reflecting functional reorganization in older adults. Overall, our findings suggest that healthy aging involves adaptive reconfiguration of neural systems, supported by experience-driven strategies that preserve semantic proficiency despite broader age-related cognitive changes.

## Introduction

Unlike most cognitive domains, semantic cognition endures relatively well into older age (Burke and MacKay, 1997; Park et al., 2002). Healthy older individuals typically maintain performance on tests assessing general knowledge, word associations, and vocabulary (Burke and Peters, 1986; Nyberg et al., 1996; Verhaeghen, 2003; Salthouse, 2004). Similarly, knowledge assessed implicitly through priming techniques exhibits constancy with age (Laver and Burke, 1993). Priming tasks reliably reveal the semantic congruency effect, where the preceding linguistic context (prime) facilitates the processing of target words that are semantically expected or related (congruent), in contrast to incongruent targets which are associated with longer reaction times. This effect remains stable across age, indicating integrity of implicit semantic processing (Laver and Burke, 1993). Building on this evidence, recent theories propose that aging involves a semanticization of cognition, reflected in greater reliance on lifelong knowledge and experience-based strategies (Zaval et al., 2015; Spreng and Turner, 2019).

Despite these well-established findings in behavioral psycholinguistics showing stable semantic cognition across age, functional neuroimaging studies have identified age-dependent changes in the neural basis of semantic processing. In electroencephalography (EEG) or magnetoencephalography (MEG) recordings, processing of meaningful stimuli is reflected in the N400 response, occurring between 200 and 600 ms after stimulus onset. The N400 is associated with a well-documented congruency effect, characterized by weaker amplitude for semantically congruent than incongruent stimuli (Kutas and Hillyard, 1980; Kutas and Federmeier, 2011). Notably, EEG studies have consistently reported a reduced N400 congruency effect in healthy aging (Kutas and Iragui, 1998; Federmeier and Kutas, 2005; Kemmotsu et al., 2012; Wlotko and Federmeier, 2012), suggesting that older adults may rely on contextual cues less than younger individuals when accessing semantic information. However, this interpretation contradicts psycholinguistic evidence.

Functional magnetic resonance imaging (fMRI) has similarly identified age-related changes in the neural organization of semantic cognition. In young adults, semantic language processing engages a predominantly left-lateralized network, including medial temporal, posterior inferior parietal, and inferior prefrontal brain regions (Binder et al., 2009). In older adults, fMRI studies have reported reduced activation in semantic-specific regions in the left hemisphere, accompanied by increased activation in the right frontal and parietal regions, particularly in the inferior prefrontal cortex (Hoffman and Morcom, 2018). This pattern of frontal and bilateral hyperactivation in older adults has been interpreted as a compensatory mechanism supporting task performance despite age-related neuronal decline (Cabeza et al., 1997).

Discrepancies between behavioral and neuroimaging findings may partly reflect methodological differences, as each approach captures different perceptual and cognitive processing stages (Payne and Silcox, 2019). Electrophysiological studies on aging have seldom examined processing stages after the N400 response. The Late Positive Complex (LPC), occurring after 500 milliseconds, has been associated with incorrect predictions based on semantic context in young adults (Van Petten and Luka, 2012). In older adults, however, the LPC has demonstrated either a reduced (Federmeier et al., 2010) or an increased (Xu et al., 2017) congruency effect. These inconsistent findings may reflect sensitivity to task-related modulations or larger individual variability in later processing stages with age.

The exact impact of aging on context processing in semantic tasks thus remains a subject of ongoing debate across disciplines (Payne and Silcox, 2019). While psycholinguistic research suggests stability in context-dependent congruency effects, imaging evidence indicates changes with age. The present study aims to bridge this gap by combining electrophysiological and behavioral measures to investigate the neural dynamics underlying lexical-semantic processing and context-driven decision-making in young (22–32 years) and old (62–69 years) adults. For this purpose, we adopt an MEG judgment task using semantically related and unrelated word pairs to probe the congruency effect, and employ a multivariate time-resolved decoding approach to characterize the neural dynamics of context processing in healthy aging, thereby advancing our knowledge of neural reorganization in older adults and informing strategies for supporting semantic function in aging populations.

## Materials and Methods

### Subjects

We studied 46 healthy right-handed (one ambidextrous, assessed using the Edinburgh Handedness Inventory) native Finnish speakers from two age groups: young adults (22 – 32 years, mean 26 years, SD 2.6) and old adults (62 – 69 years, mean 65 years, SD 1.6). All subjects (25 in the young group [12 females], 21 in the old group [13 females]) were screened for pre-existing neurological disorders, learning disabilities, language disorders and cognitive impairment (Montreal Cognitive Assessment score 26 or above), had normal or corrected-to-normal vision and were not considered part of Covid-19 risk groups due to the pandemic situation at the time of the recordings. They signed an informed consent before participating and were compensated for lost working hours. The ethical statement for the study was obtained from the Aalto University Research Ethics Committee.

### Neuropsychological assessments

A series of neuropsychological tests were employed to examine potential age-related variances in subjects’ verbal and non-verbal abilities. From the fourth edition of the Wechsler Adult Intelligence Scale, WAIS-IV (Wechsler, 2008), four subtests were applied: Block Design (visuoconstructive reasoning), Similarities (abstract verbal reasoning, semantic knowledge), Vocabulary (word knowledge, verbal concept formation), and Digit-Symbol-Coding (perceptive-motor processing speed, working memory). In addition, we used the Trail Making Test (TMT) A (attention, processing speed) and B (attention, set-shifting, cognitive flexibility) (Reitan, 1958). To address verbal skills, two verbal fluency tests were incorporated, where subjects were to produce as many words as possible within 60 seconds for a given category. A semantic (animals) and a phonetic (words beginning with letter s) test were included (Benton, 1968). Raw scores were converted to a common scale using sample-based Z-scores. In addition, age-adjusted standardized Z-scores were derived based on Finnish normative data. Differences in raw and standardized neuropsychological test scores between age groups were assessed with two-tailed independent-samples t-tests.

### Experimental design

We recorded MEG data from participants while they were performing a lexical-semantic judgment task. During the task, written Finnish word pairs were projected onto a screen placed at 120-cm distance in front of the participants using the Presentation software (version 22.0; Neurobehavioral Systems Inc.). The words were presented one at a time. The word pairs consisted of a prime and a target word, and the subjects were asked to indicate whether the words were semantically related/congruent (e.g., banana – orange) or unrelated/incongruent (e.g., banana – police) by responding either with their right index or middle finger. The prime and target words were presented sequentially, such that each word was shown for 300 ms, with a 700 ms interval between the words. The subjects were given 2500 ms time to respond, during which time a blank screen was presented, and an additional 1000 ms (500 ms fixation cross + 500 ms blank screen) to prepare for the next word pair. The total inter-stimulus-interval between the pairs was then 3500 ms. The responses were registered using an optical device, and response fingers and corresponding stimulus conditions were counterbalanced between the subjects within each age group. A short practice task was introduced before the recordings. In total, 108 congruent and 108 incongruent word pairs were presented to the participants in a random order. Structural magnetic resonance (MR) images were obtained in a separate session after the MEG measurement.

### Stimuli

For the prime words, 216 common Finnish nouns were selected (mean word length [len] = 5.78 letters, mean word frequency per million [wpm] = 49 015 [The Finnish Internet Parsebank, Kanerva et al., 2014]). Each prime word was assigned both a semantically congruent and a semantically incongruent target word (len = 5.86, wpm = 42 056). Semantic congruency between the prime and target words was assessed using a distributional model of the semantic system, word2vec, trained with a lemmatized 126 617 892 - word Finnish Internet Parsebank (Kanerva et al., 2014). Word2vec is a computational method for extracting distributional vector space representations of a lexicon which can be used to derive semantic similarities between words using cosine similarity (Mikolov et al., 2013). The average cosine similarity for congruent pairs was.6675 (SD =.1702), while for incongruent pairs, it was.0398 (SD =.06612).

Two stimulus sets, A and B, sharing the same set of primes were crafted. In set A, half of the prime words were paired with a congruent target and the other half with incongruent targets. In set B, the same prime stimuli were used in the other condition, i.e., the first half was paired with incongruent and the latter half with congruent targets. Each participant was presented with 216 unique primes paired with targets drawn either from set A or set B, counterbalanced between age groups. The stimulus subsets were matched for word length and word frequency (*p* <.05, independent samples t-test).

### Analysis of behavioral results

In the lexical-semantic judgment task, participants were required to actively categorize the target stimulus as either congruent (semantically related) or incongruent (unrelated) and give their manual response by lifting their right index or middle finger (counterbalanced between age groups). Differences in response times and accuracies between age groups and stimulus conditions were studied with two-tailed independent-samples t-tests. Response accuracy was determined as the ratio between correct and incorrect responses.

### MEG and MRI recordings

MEG data were recorded at Aalto University MEG Core using a 306-channel (204 gradiometers and 102 magnetometers) Vectorview device (MEGIN Oy, Helsinki, Finland) inside a high-end 3-layer magnetically shielded room (Imedco AG, Hägendorf, Switzerland). MEG signals were acquired at a sampling frequency of 1000 Hz and filtered to 0.1 – 330 Hz during acquisition. Vertical and horizontal electro-oculogram (EOG) signals were recorded to identify eye blinks and saccades. Five head position indicator (HPI) coils were affixed to the head, and their positions relative to anatomical landmarks (nasion and preauricular points) were digitized using an electromagnetic tracking system (Fastrak, Polhemus Inc., Colchester, VT, USA). Continuous head position indication (cHPI) was used to compensate for head movements in relation to the sensor array during the measurement. Structural MR images were acquired at Aalto University Advanced Magnetic Imaging (AMI) Centre using a 3 T Magnetom Skyra whole-body scanner (Siemens Healthcare, Erlangen, Germany) and a standard T1-weighted Magnetization Prepared Rapid Gradient Echo (MPRAGE) sequence.

### MEG preprocessing

MEG data were preprocessed using MaxFilter version 2.2.15 (MEGIN Oy) and MNE-Python version 1.5.1 (Gramfort et al., 2014). Temporally extended signal space separation (tSSS; Taulu and Simola, 2006) was applied to suppress signal artifacts originating from outside the brain. Data from different subjects and measurements were transformed into a common head position enabling sensor-level analysis at the group level. Independent component analysis (ICA) using the Extended Infomax algorithm (Lee et al., 1999) and a 30-component decomposition was used to suppress ocular, cardiac, and other physiological artifact components. On average, 3.3 (SD = 2.5) components per subject were removed. The preprocessed MEG data were low-pass filtered at 40 Hz and segmented to event-related epochs from 200 ms before to 1000 ms after the stimulus onset. Baseline correction was applied using the 200 ms interval preceding the stimulus. Epochs with amplitudes exceeding 3000 fT/cm for gradiometers or 4000 fT for magnetometers were discarded (0.44% of all epochs) and the number of epochs per condition were equalized for each subject.

### Evoked activity and source modelling

MEG evoked responses were obtained by averaging across epochs for each condition. For source-level visualization, the cortical-level spatiotemporal distribution of the evoked activity was obtained through cortically constrained L2-minimum-norm estimates (Hämäläinen and Ilmoniemi, 1994) using MNE-Python version 1.5.1 (Gramfort et al., 2013). Sources perpendicular to the cortical surface were favored using a loose orientation constraint of.2, and a depth weighting of.8 was used to reduce the bias towards superficial sources. Noise covariance matrices were computed from the 200 ms pre-stimulus baselines of the prime epochs (the initial words in a pair), and noise normalization was applied to obtain dynamic statistical parametric maps (dSPM, Dale et al., 2000). Cortical surfaces needed for anatomically informed source localization were derived from the individual MR images using Freesurfer 6. 0 (Fischl, 2012). Co-registration between structural images and MEG data was performed using digitized anatomical landmarks (nasion and pre-auricular points), coil positions, and additional reference points from the scalp. For each subject, a head conductor model was constructed using a single-layer boundary-element model (BEM) from the inner skull surface consisting of 20484 triangles. A source space with 10242 vertices (ico-5) per hemisphere was employed in the forward model computation. For group-level analyses, the source estimates were morphed onto a template brain (FreeSurfer fsaverage). Anatomical parcellations (Desikan et al., 2006; Glasser et al., 2016) were used to segment the cortical surface into regions of interest (ROI).

### MEG analysis

#### Difference in evoked activity between conditions

To detect alterations in evoked activity associated with semantic congruency, we first studied the difference between congruent and incongruent conditions at the sensor level. The semantic congruency effect was determined as the point-by-point difference in gradiometer activation by subtracting the evoked responses in the congruent condition from those in the incongruent condition. Each sensor in a gradiometer pair was handled independently. The significance of the congruency effect was studied in three designated time windows (200 – 400 ms, 400 – 600 ms, and 600 – 800 ms) using two-tailed one-sample t-tests (H0 = 0). The hypotheses were tested independently for four averaged sensor selections (left temporal, right temporal, left frontal, and right frontal) for each age group. To account for multiple comparisons, the p-values were adjusted using the Benjamini–Hochberg procedure (false discovery rate, FDR).

#### Region of interest analysis

Age-related differences in the source-level congruency effects were examined by comparing activation differences between age groups within two regions of interest (ROIs). For each participant, the semantic congruency effect was computed as the point-by-point difference between the stimulus conditions. The first ROI was defined in the left superior and middle temporal cortex (auditory label; Glasser et al., 2016), corresponding to the typical neural correlates of the N400 (Helenius et al., 1998; Lau et al., 2008; Kutas and Federmeier, 2011), and the second ROI covered the left frontal cortex, where the LPC is typically observed (Van Petten and Luka, 2012). Mean activation time courses within these ROIs were derived using the PCA flip method, which applies singular value decomposition to extract the dominant component and implements scaling and sign-flipping to ensure consistent orientation across source estimates. Group differences were assessed within three time windows (200–400 ms, 400–600 ms, and 600–800 ms) using one-tailed independent-samples t-tests, with FDR correction applied to control for multiple comparisons.

#### Laterality Index and Temporal-Frontal Index

To study age-related differences in activation distribution across the hemispheres, we used Laterality Index (LI) which has been used in previous studies to determine language lateralization (Doss et al., 2009; Tanaka et al., 2013; Raghavan et al., 2017). Following the approach of Tanaka et al. (2013) and Raghavan et al. (2017), we defined subject-specific activation thresholds as 50% of the maximum value over both hemispheres within each of the three selected time windows (200–400, 400–600, and 600–800 ms). LIs were defined as:

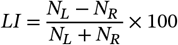

where *N*_*L*_ and *N*_*R*_ represent the number of above-threshold vertices in the left and right hemispheres. The LI values range between −100 and 100, and LI ≥ 10 was considered left hemisphere dominance, LI ≤−10 right hemisphere dominance, and LI between −10 and 10 bilateral (Doss et al., 2009; Tanaka et al., 2013; Raghavan et al., 2017).

Following a similar approach, we determined Temporal-Frontal Indices (TFI) to investigate the age-related differences in activation distribution between temporal and frontal brain regions. Here, we used the Desikan-Killiany parcellation (Desikan et al., 2006) to construct two bilateral ROIs covering the entire temporal and frontal cortices in both hemispheres. The activation thresholds were defined as 50% of the maximum value over both ROIs within each time window. TFIs were defined as:

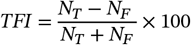

where *N*_*T*_ and *N*_*F*_ represent the number of above-threshold vertices in the temporal and frontal ROIs. Ranging between −100 and 100 the more positive values correspond with temporal-over-frontal emphasis, and the more negative values with frontal-over-temporal emphasis. Since the vertex count between the ROIs is not equal, this index is not symmetrical, unlike the LI, but it describes the relative ratio between temporal and frontal activation.

#### Temporal decoding of the semantic congruency

A time-resolved within-subject decoding analysis was conducted to investigate the temporal dynamics of the semantic congruency effect. The analysis was performed using MNE-Python (version 1.5.1; Gramfort et al., 2014) and scikit-learn (version 1.3.1; Pedregosa et al., 2011). Logistic regression classifiers were employed with stratified 10-fold cross-validation, and classifier performance was evaluated using the Area Under the Receiver Operating Characteristic Curve (ROC-AUC) scores. Binary classification between congruent and incongruent epochs was repeated at each time point, generating a time-series of ROC-AUC scores describing the evolution of classification performance over time. Group-level results were obtained by averaging within-subject results. To determine whether decoding scores deviated significantly from chance level, cluster-based permutation tests were conducted with 1000 permutations and a cluster-forming threshold of *p* <.05.

To examine the spatial origin of the semantic congruency effect, classifier weights for each subject were transformed into activation patterns using the method proposed by Haufe et al. (2014). The neural generators of these reconstructed patterns were localized using the inverse solutions described earlier. The resulting relevance patterns indicate the extent to which different cortical regions contain condition-specific information about the stimuli. Prior to group-level averaging, individual relevance patterns were morphed onto the fsaverage template brain.

Furthermore, Pearson’s correlation analysis was conducted to evaluate the relationship between decodability and behavioral performance. Decodability scores were averaged across 200 – 800 ms, and the correlations between the resulting mean scores and behavioral reaction times were analyzed.

#### Temporal decoding of the age group

A similar time-resolved decoding approach was employed to directly assess differences in the semantic congruency effect between the young and the old age groups. Here, logistic regression classifiers were trained on the evoked incongruent-congruent contrast responses labeled by age group. To mitigate overfitting, a group shuffle-split cross-validation scheme was implemented, adhering to subject boundaries and ensuring that the test set did not incorporate samples from participants present in the training set. In practice, the data was shuffled and randomly divided into training and test sets, with six subjects included in each test split. This re-shuffling and splitting process was repeated 50 times. Age groups were initially split separately from each other and subsequently combined to ensure equal representation in both the training and test sets. As in the previous analysis, classifier weights were used to compute cross-group relevance patterns (Haufe et al., 2014). Source reconstruction was performed at the subject-level following the previously described steps, and the final cross-group cortical relevance patterns were obtained by averaging these source estimates.

## Results

### Neuropsychological assessments

The general cognitive performance of the participants in the two age groups, assessed using a neuropsychological test battery, is described in **Figure 1**. The old group scored lower in perceptual reasoning and processing speed tests (Trail Making Test A [*p* =.002], Trail Making Test B [*p* =.003], WAIS-IV Block design [*p* <.001], WAIS-IV Digit-Symbol-Coding [*p* =.001]), whereas no age-related differences were found in verbal tests (WAIS-IV Similarities [*p* =.283], WAIS-IV Vocabulary [*p* =.287], Phonemic Verbal Fluency [*p* =.663], Semantic Verbal Fluency [*p* =.797]) (**Figure 1** upper section). When adjusted for age, both groups exhibited slightly higher performance than their respective age-matched reference populations (**Figure 1** lower section). However, this advantage was more pronounced among older adults than the young in Trail Making Test A (*p* <.001), Trail Making Test B (*p* =.046), WAIS-IV Digit-Symbol Coding (*p* =.001), and Phonemic Verbal Fluency test (*p* =.048).

**Figure 1.**
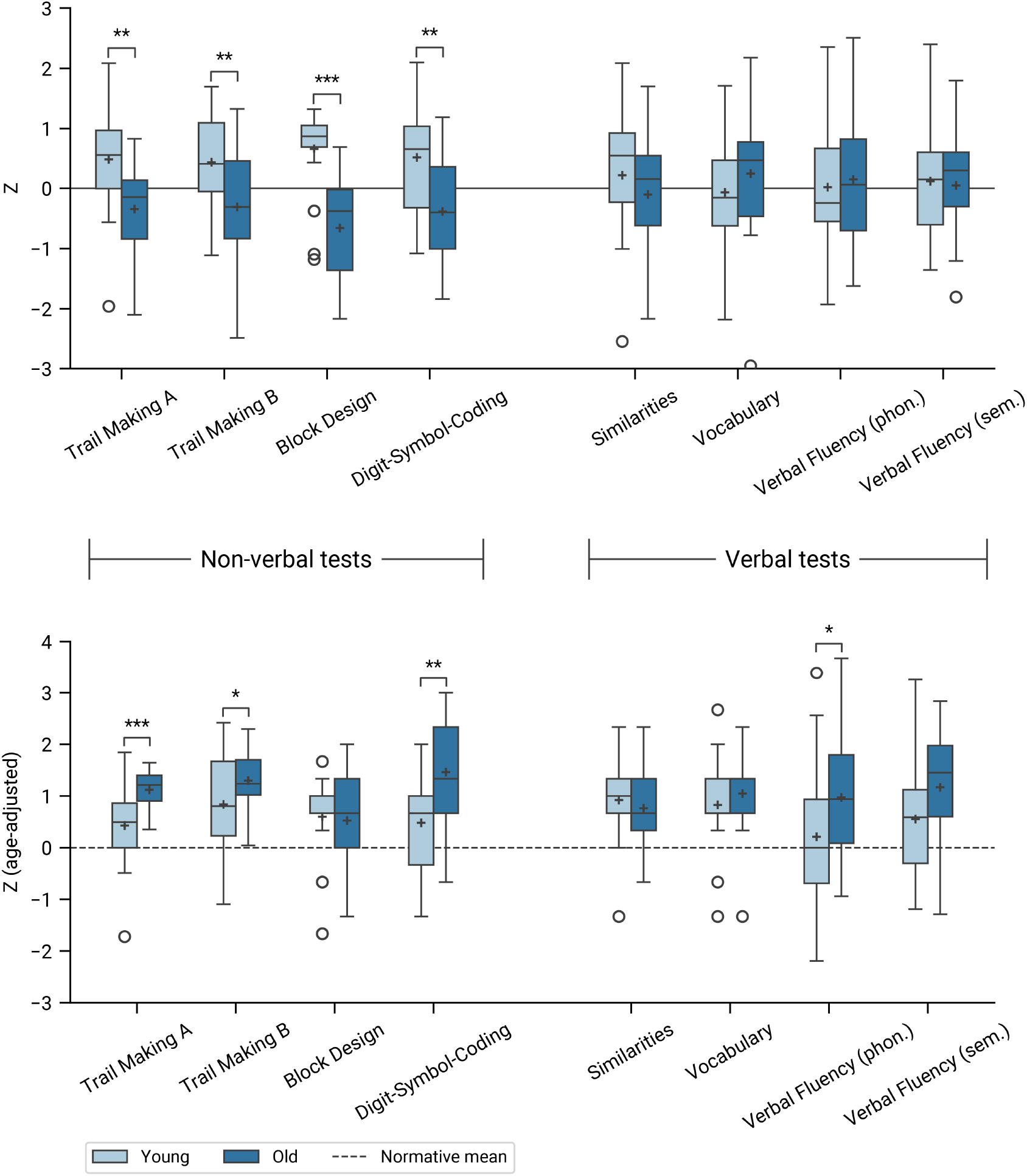
Neuropsychological assessment results. The upper section presents raw sample-based Z-scores, while the lower section displays Z-scores standardized using age-matched normative data. Significant differences between age groups are highlighted (independent-samples t-tests).

### Behavioral results

During the MEG recordings, the participants performed a lexical-semantic judgment task during which they were required to actively categorize the target stimulus as either congruent (semantically related) or incongruent (unrelated). The performance in the task for young and old age groups, with respect to response times and accuracy, is presented in **Figure 2**. The old group had significantly faster response times in both the congruent and incongruent conditions (mean 809.4 ms, SD 235.9) than the young (mean 916.3 ms, SD 228.4, *p* <.001). On average, the gap between age groups was 106.9 ms with no discernible difference between conditions. On the other hand, response times for incongruent stimuli were significantly slower than for congruent stimuli in both young (congruent: mean 894.6 ms, SD 287.1, incongruent: 938.0 ms, SD 289.7) and old age groups (congruent: mean 788.0 ms, SD 241.5, incongruent: mean 830.7 ms, SD 230.4, *p* <.001). On average, the difference between conditions was 43.1 ms. Here, no age-related variation was observed.

**Figure 2.**
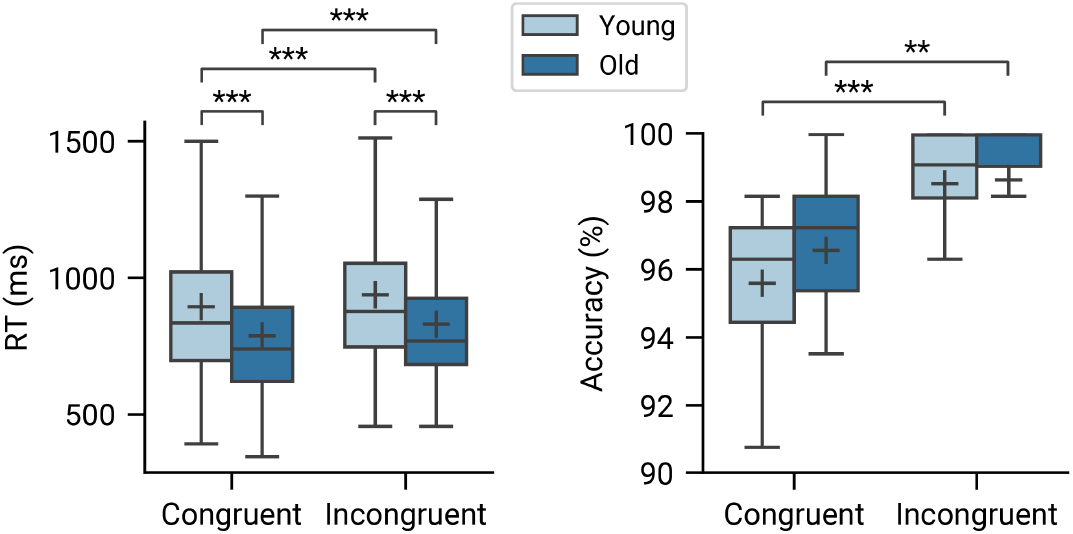
Behavioral response times and accuracy rates (defined as the ratio of correct to incorrect responses). Comparisons between age groups and stimulus conditions were conducted using independent samples t-tests. *** *p* <.001, ** *p* <.01, * *p* <.05

Response accuracy was determined by the ratio between correct and incorrect responses. Correct responses exceeded 95% across both age groups and conditions, and no differences in accuracies were found between the groups. Accuracies for incongruent stimuli were marginally greater than those for congruent stimuli in both young (congruent: mean 95.6%, SD 2.3, incongruent: mean 98.5%, SD 2.1, *p* <.001) and old (congruent: mean 96.6%, SD 2.5, incongruent: mean 98.6%, SD 2.1, *p* =.006) participants.

### Age-related differences in semantic MEG responses and their neural correlates

We examined variations in MEG responses associated with the semantic judgment task across the two age groups, with a particular focus on the semantic congruency effect, defined as the difference in neural responses between incongruent and congruent conditions.

Significant congruency effects in sensor-level MEG evoked responses (*p* <.05, two-tailed one-sample t-test, FDR-corrected) were observed in both age groups within the 200 – 800 ms window following stimulus onset (**Figure 3**). In the young group, the effect was prominent in all the time windows at the left temporal (200 – 400 ms [*p* =.001], 400 – 600 ms [*p* <.001], 600 – 800 ms [*p* =.013]), and left frontal (200 – 400 ms [*p* =.002], 400 – 600 ms [*p* =.003], 600 –800 ms [*p* =.002]) sensors, and at the right frontal sensors in the 400 – 600 ms time window (*p* =.031). In the old group, a congruency effect was observed at the left temporal sensors only in the 400 – 600 ms time window (*p* =.011) and in all the time windows at the left frontal sensors (200 – 400 ms [*p* =.002], 400 – 600 [*p* <.001], 600 – 800 ms [*p* =.031]) and right frontal sensors (400 – 600 ms [*p* =.002]). Additionally, a subtler congruency effect was detected at the right temporal sensors in both the young (200 – 400 ms [*p* =.030], 400 – 600 ms [*p* =.022]) and old groups (200 – 400 ms [*p* =.020]).

**Figure 3.**
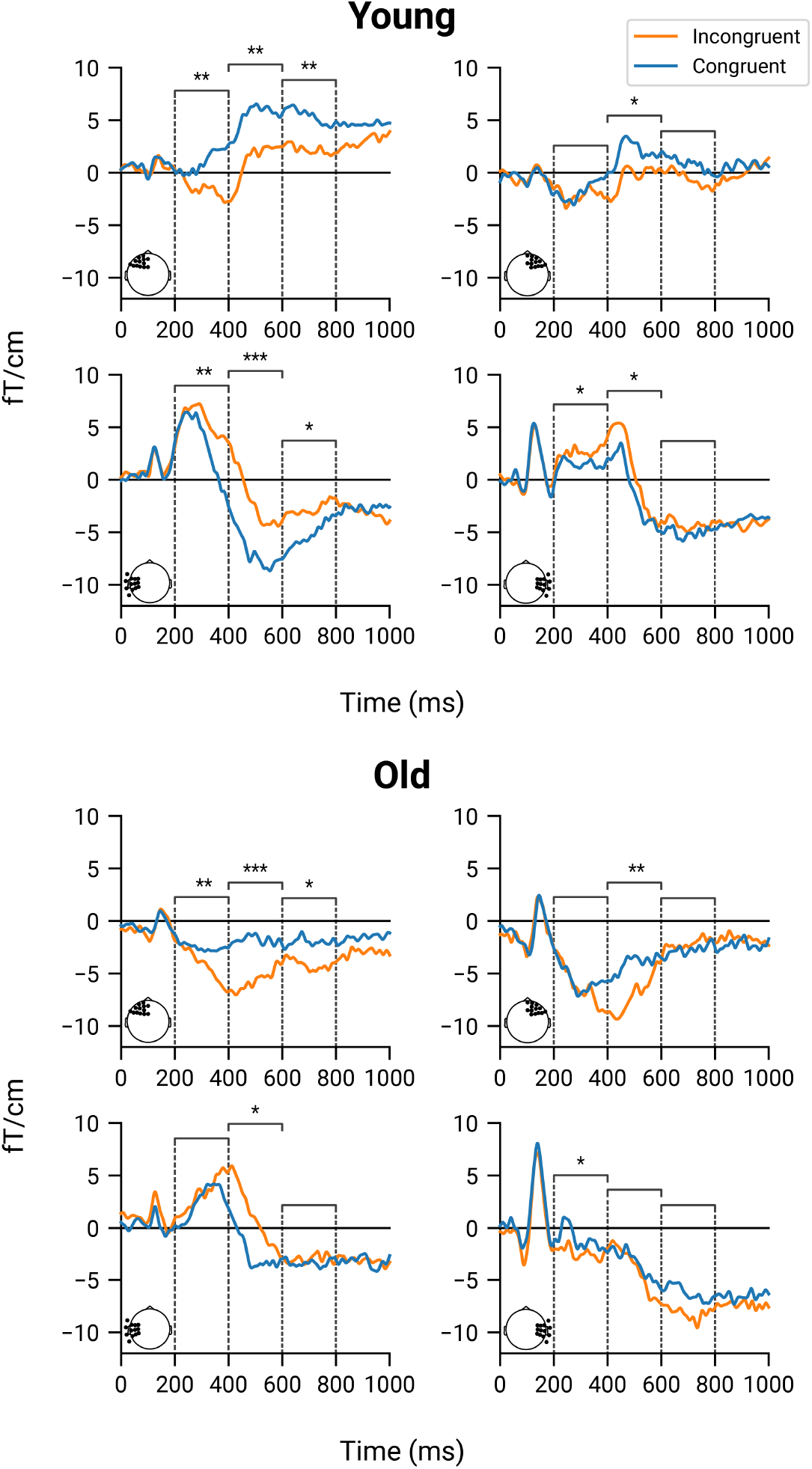
Evoked responses by age group over four sensor selections: left-frontal, right-frontal, left-temporal, and right-temporal. The significance of the congruency effect between conditions was tested in three time windows: 200–400 ms, 400–600 ms, and 600–800ms (two-tailed one-sample t-test, *** *p* <.001, ** *p* <.01, * *p* <.05, FDR-corrected).

In the young group, a characteristic N400 effect (incongruent > congruent), originating from the left superior temporal and middle temporal cortices, was observed between 300 and 400 ms (**Figure 4A**). Similar neural correlates have previously been associated with the N400 response in young individuals (Helenius et al., 1998; Lau et al., 2008; Kutas and Federmeier, 2011). In the old group, the N400 response peaked slightly later, just after 400 ms, in the left temporal sensors (**Figure 3**). No distinct source-level equivalent of the N400 effect (incongruent > congruent) was identified for the old participants, and the congruency effect exhibited significant age-related attenuation during the 200 – 400 ms interval in the left superior and middle temporal ROI (T = 2.74, *p* =.013, FDR-corrected). Instead, the N400 response in the old group was associated with widespread activation in left inferior frontal, middle frontal, and posterior temporal regions, as well as right superior temporal, middle temporal, and middle frontal regions (**Figure 4A**).

**Figure 4.**
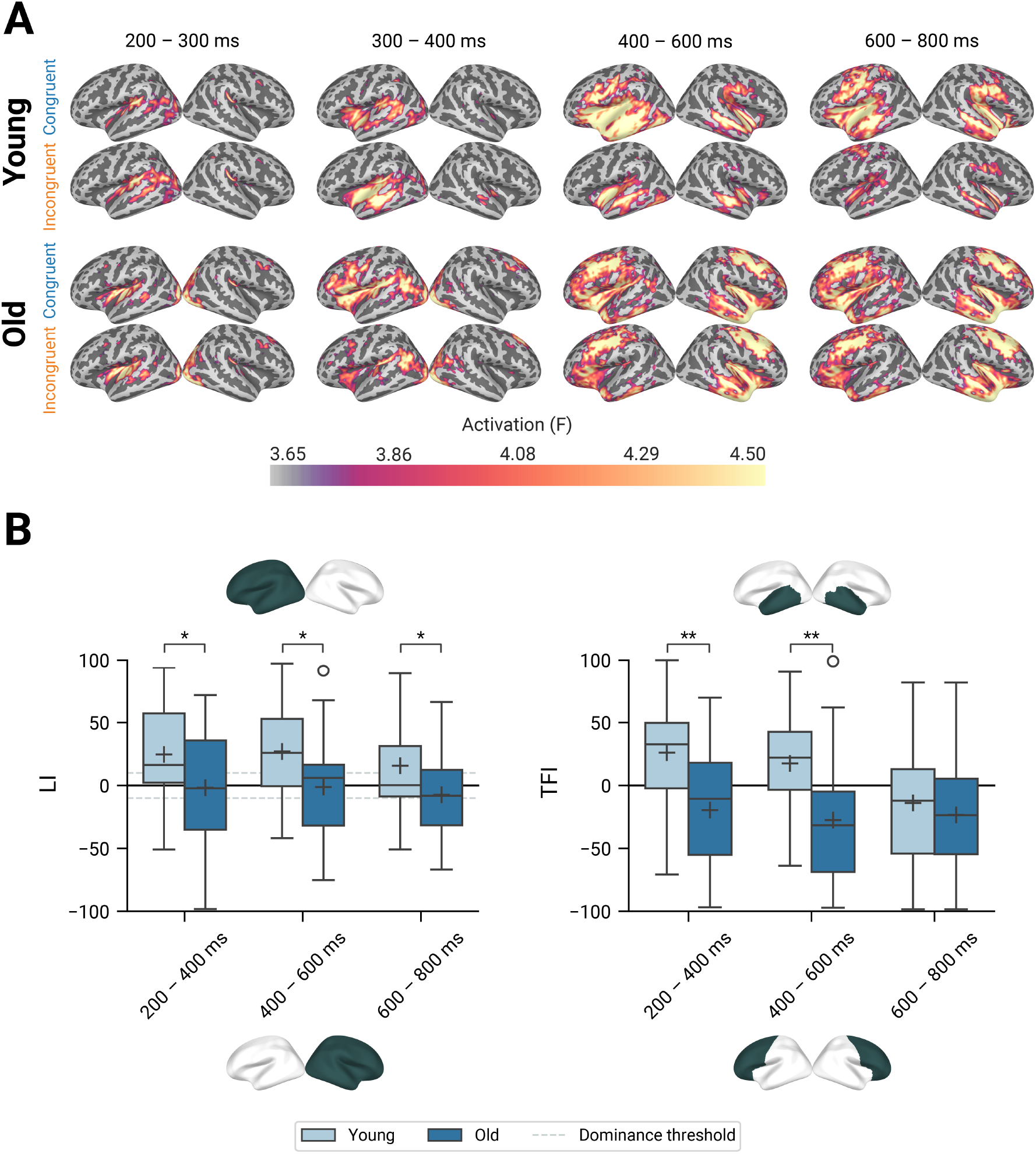
Minimum-norm estimates. **A**. Cortical source estimates (dSPM) of congruent and incongruent word pairs. **B**. Differences in Laterality Indices (LI) and Temporal-Frontal Indices (TFI) between age groups tested in three time windows (two-tailed independent samples t-test, *** *p* <.001, ** *p* <.01, * *p* <.05, FDR-corrected). LI > 0: left hemisphere emphasis; LI < 0: right hemisphere emphasis. TFI > 0: temporal-over-frontal activity; TFI < 0: frontal-over-temporal activity. The dashed line refers to the threshold of hemispheric dominance (LI ≥ 10: left hemisphere dominance, LI **≤**−10: right hemisphere dominance [Doss et al., 2009; Tanaka et al., 2013; Raghavan et al., 2017]).

In the young group, a later, inverted congruency effect (congruent > incongruent) was observed from 400 to 800 ms (**Figure 3**). The neural correlates of this LPC-like effect extended from the left temporal to the left middle frontal and inferior frontal regions (**Figure 4A**). In the old group, activation patterns associated with the LPC response were observed both in the left and right middle frontal and inferior frontal regions, as well as in the right middle temporal regions (**Figure 4A**). However, older participants demonstrated significantly weaker congruency effects than the young during the 400 – 600 ms interval in the left frontal ROI (T = 2.22, *p* =.032, FDR-corrected) and the 600 – 800 ms interval in the left superior and middle temporal ROI (T = 2.91, *p* =.013, FDR-corrected).

To assess the overall distribution of cortical activation during the task, Laterality Indices (LI) and Temporal-Frontal Indices (TFI) were computed for selected time windows (**Figure 4B**). The young group exhibited significantly greater left-hemisphere-dominant activation (LI ≥10) across all time windows compared to the old group (*p* =.049, FDR-corrected), whose activation patterns were generally more bilateral (LIs between −10 and 10). Furthermore, in examining the relative balance of activation between temporal and frontal regions, the young participants demonstrated significantly greater emphasis on the temporal cortex (positive TFIs) during the 200 – 600 ms time window (*p* =.003, FDR-corrected), whereas in the old group, activation was significantly more frontally distributed (negative TFIs). In the later processing stage (600 – 800 ms), activation distribution in the young group shifted towards frontal regions, and no TFI differences between age groups were observed (*p* =.519, FDR-corrected).

### Temporal decoding

#### Decoding the stimulus condition

We examined the temporal evolution of the semantic congruency effect by training a series of logistic regression classifiers to differentiate between congruent and incongruent evoked responses at each time point within each participant. Classifier performance was subsequently evaluated and plotted as a function of time (**Figure 5A**). Stimulus conditions became decodable approximately 300 ms after stimulus onset, with peak decoding performance occurring shortly after 400 ms. At this peak, ROC-AUC scores reached.6 in the young cohort and.7 in the old cohort. A gradual decline in decoding performance was observed in the older group after 500 ms, whereas performance in the younger group remained relatively stable between 400 and 800 ms.

**Figure 5.**
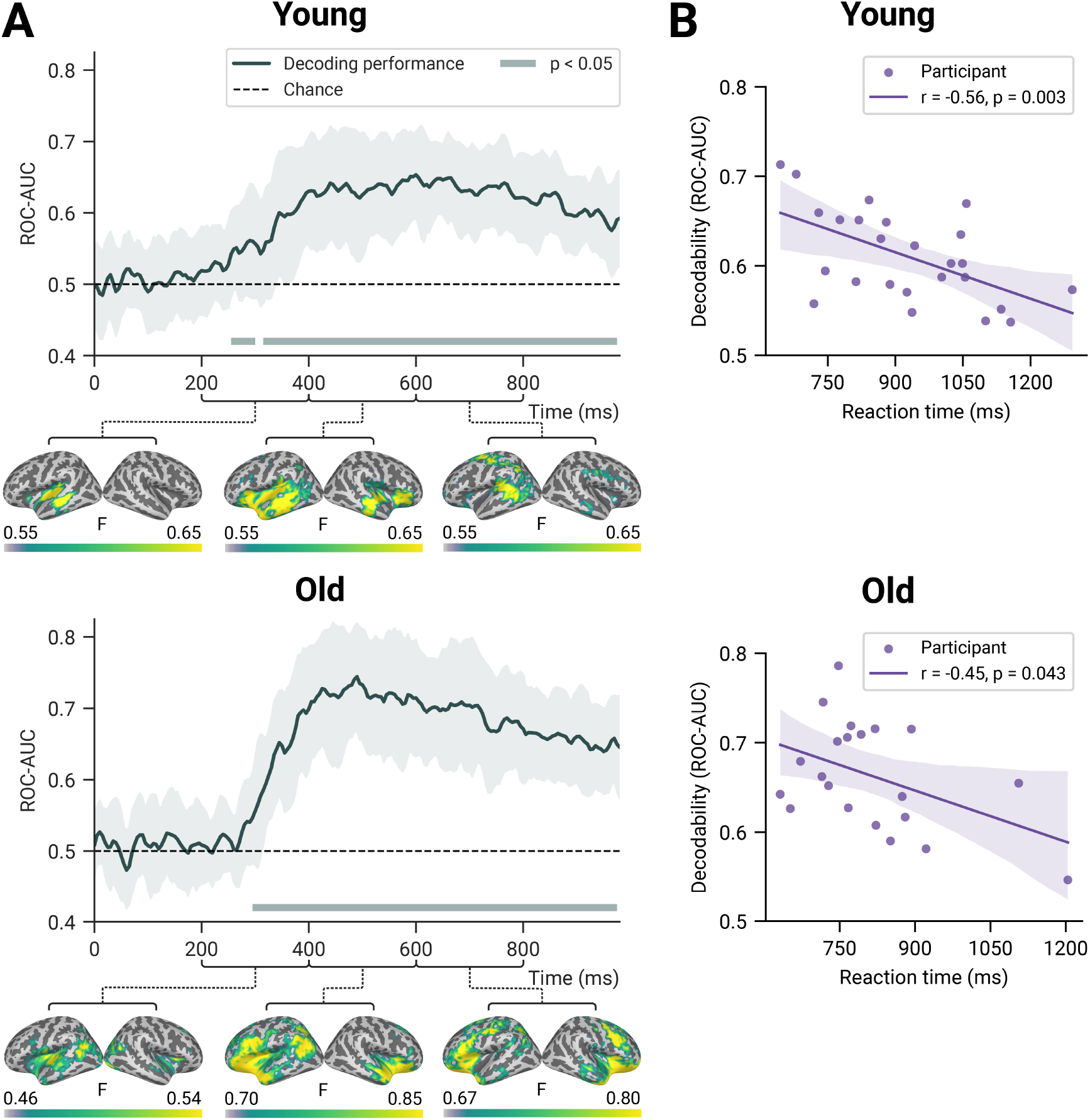
Decodability of stimulus condition (congruent vs. incongruent). **A**. Time-resolved decoder performance (+/- SD), quantified by the area under the receiver operating characteristic curve (ROC-AUC), is presented. Gray bars indicate time intervals during which decoding accuracy significantly exceeded chance level (*p* <.05, two-tailed, within-group cluster-based permutation tests). Corresponding cortical relevance patterns (Haufe et al., 2014) highlight the neural sources contributing to classifier performance based on MEG signal features. **B**. Correlation between decodability (averaged from 200 to 800 ms) and behavioral reaction times (Pearson’s r +/- SD).

Cortical relevance patterns were derived by transforming classifier weights into activation patterns and projecting them onto the cortical surface. The activation patterns presented in **Figure 5A** highlight the brain regions most relevant for congruent-incongruent classification within specific time windows, separately for both age groups.

In the young group, the decoder performance between 200 and 400 ms was predominantly associated with the left middle and superior temporal regions, corresponding to the neural correlates of the N400 effect observed (**Figure 4A**). In the old group, similar left temporal patterns emerged, albeit with a more posterior distribution (**Figure 5A**). During the 400 – 600 ms window, these relevance patterns shifted toward the left frontal regions, as well as the right anterior temporal and inferior frontal regions, similarly to the cortical source estimates (**Figure 4A**).

In the young group, post-N400 relevance patterns in the 400 – 600 ms window extended to the left inferior frontal and inferior temporal regions, as well as right middle temporal and inferior frontal regions. Subsequently, between 600 – 800 ms, a pronounced left posterior temporal emphasis was observed. Compared to the LPC source estimates for the young group in **Figure 4A**, these relevance patterns exhibited greater temporal and reduced frontal distribution. In the old group, the LPC relevance patterns between 600 and 800 ms were associated with left and right middle frontal and inferior frontal, as well as right middle temporal regions (**Figure 5A**), corresponding to the LPC source estimate results.

The relationship between MEG response decodability (averaged from 200 to 800 ms) and behavioral reaction times (averaged across conditions) was examined using Pearson’s correlation analysis. Higher decodability scores were associated with faster reaction times in both age groups (**Figure 5B**). Both younger and older participants showed a significant negative correlation, such that the better the stimulus condition decodability between 200 – 800 ms was for a given individual, the faster was their response time.

#### Decoding the age group

Age-related differences in the semantic congruency effect were examined using a similar temporal decoding approach. Logistic regression models were trained to classify between the young and old age groups at distinct time points based on the evoked contrast responses (incongruent minus congruent). Age group classification was successful within the 200–800 ms time window, with ROC-AUC scores peaking at.8 (**Figure 6**).

**Figure 6.**
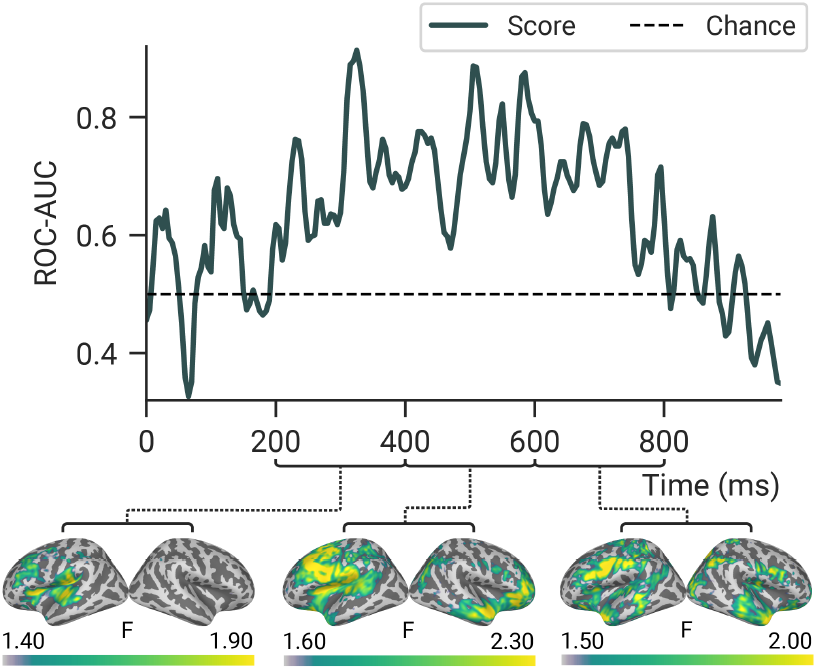
Decodability of age group, based on the semantic congruency effect in the MEG responsess. Time-resolved decoder performance (ROC-AUC) over time is shown for the classification between younger and older adults based on the semantic congruency effect (incongruent minus congruent). Corresponding cortical relevance patterns (Haufe et al., 2014) indicate the neural sources contributing to classification performance.

The cortical relevance patterns in **Figure 6** highlight several brain regions that contributed to the decoding performance, thereby reflecting functional age-related differences in the semantic task. Within the 200 – 400 ms window, the primary contributing regions included the left middle temporal, superior temporal, and inferior frontal cortices. Between 400 and 800 ms, a prominent shift was observed toward the left middle frontal, as well as right anterior temporal and inferior frontal regions. Finally, in the 600 – 800 ms window, decoder performance was predominantly associated with age-related activation differences in the left middle frontal cortex.

## Discussion

This study examined the effects of healthy aging on the neural underpinnings of semantic congruency judgments by integrating MEG and behavioral measures in younger and older adults. While behavioral performance remained stable across age groups, age-related differences emerged in the time-resolved neuronal responses. Multivariate MEG decoding further revealed that greater neural separation between semantic concepts facilitated faster decision-making. These findings suggest an adaptive reorganization of neural systems preserving semantic proficiency in aging.

### Preserved behavioral performance despite age-related changes in semantic MEG responses

Both age groups completed the lexical-semantic judgment task with high accuracy, exceeding 95% correct responses. Older adults in their sixties, however, demonstrated significantly faster response times than their younger counterparts, indicating that advancing age was not associated with impaired task performance. On the contrary, healthy and motivated older adults may even outperform young adults. These findings are consistent with prior psycholinguistic research indicating that semantic cognition remains largely stable across the lifespan (Burke and MacKay, 1997; Park et al., 2002). The neuropsychological assessments further supported this conclusion, showing comparable verbal performance across age groups, but superior non-verbal reasoning and processing speed in younger adults.

The observation that older adults exhibited faster response times than younger adults in the semantic judgment task was somewhat unexpected, given that processing speed generally declines by age (Salthouse, 1996). However, older adults’ greater knowledge reserve, accumulated through life experience, may facilitate more efficient decision-making. Older adults are likely to favor automatic strategies grounded in experience-based knowledge and semantic networks acquired over years, which are largely preserved beyond age 60 (Li et al., 2004; Salthouse, 2004; Zaval et al., 2015; Spreng and Turner, 2019). This semanticization of cognition may underlie their faster response times, reflecting a strategic shift toward rapid, knowledge-based processing over slower, deliberative analysis. In addition, higher motivation and cognitive aptitude (age-adjusted neuropsychological test scores exceeding the young) may also have contributed to the superior performance in older participants. Regardless of the specific explanatory mechanisms, these findings underscore that older adults can maintain high proficiency in processing semantic relationships and may even surpass younger adults in terms of response speed.

Despite preserved behavioral performance, age-related alterations in MEG responses were observed. As anticipated, young adults demonstrated an N400 response peaking between 300 and 400 ms in the temporal cortex. In the old group, the N400 peaked slightly later, after 400 ms, and lacked a specific neural correlate, instead showing widespread activation in the left frontal and posterior temporal regions. Furthermore, the characteristic N400 congruency effect, marked by greater activation for incongruent versus congruent stimuli, was more prominent in the young group than the old, consistent with prior research (Kutas and Iragui, 1998; Federmeier and Kutas, 2005; Kemmotsu et al., 2012; Wlotko and Federmeier, 2012). Despite these differences, decoding analyses showed a sharp increase in classification accuracy from 300 ms onward in both age groups, indicating preserved sensitivity to congruency with age. Thus, the attenuated N400 effect in older adults likely reflects broader neural reorganization of the semantic network rather than diminished contextual utilization or semantic deficits.

Following the N400, a later, frontotemporal LPC-like response emerged between 400 – 800 ms in the young group, showing a clear congruency effect opposite to the N400 effect (congruent > incongruent). This response was less pronounced in older adults, with no significant amplitude difference between conditions. Nonetheless, decoding analysis detected congruency effects in both groups. As the LPC is considered to reflect more intentional processes such as decision-making and explicit semantic access (Hoshino and Thierry, 2012; Yang et al., 2019), its prominence in young adults may reflect greater reliance on deliberative, resource-intensive decision-making strategies.

### Frontal shift and increased bilaterality in older adults reflect neural reorganization and the semanticization of cognition

Overall, a distinct activation distribution across cortical regions was observed between younger and older adults, characterized by a shift from temporal to frontal regions and from left-lateralized to more bilateral processing in older adults. Age-related differences were most pronounced between 200 and 600 ms after stimulus onset. These findings align with previous fMRI findings on posterior–anterior shifts and reduced hemispheric asymmetry in aging (Cabeza, 2002; Davis et al., 2008; Hoffman and Morcom, 2018). In the present study, we were able to characterize the temporal aspects of these age-related shifts using MEG.

This neural reorganization, likely reflecting the context-dependent engagement of a broad network of brain regions, may indicate semanticization of cognition and reliance on accumulated experience in aging (Spreng and Turner, 2019). We argue that increased frontal recruitment and bilaterality in the older adults represent not just compensation for age-related decline (Cabeza et al., 1997), but functionally adaptive changes in the semantic system. Beyond core semantic processing, task demands likely influenced the later-stage neural patterns. Frontal regions, critical for semantic control and decision-making (Badre and Wagner, 2007; Jackson, 2021), were engaged in both groups after 600 ms, likely reflecting active decision-making associated with the task and involvement of higher-order cognitive strategies for resolving semantic incongruities.

### Distinctive neural representations support semantic judgment

We applied multivariate time-resolved decoding to capture age-related discrepancies in the temporal dynamics of lexical-semantic processing at the individual level. Unlike traditional univariate methods, which may obscure subtle neural variations due to signal averaging, multivariate approaches analyze activation patterns across sensors, increasing sensitivity (Haynes and Rees, 2006; Grootswagers et al., 2017). This can provide more detailed insights into age-related differences underlying semantic congruency evaluations (King and Dehaene, 2014; Contini et al., 2017).

As discussed, within-group decoding was highly sensitive to congruency effects, and the associated cortical relevance patterns demonstrated enhanced frontal and bilateral recruitment in older participants, paralleling source modelling results. Notably, decoding performance was significantly correlated with task performance: faster responses were associated with higher decoding scores across both age groups. This suggests that the neural differentiation between congruent and incongruent words, reflected in high decoding performance, modulates the efficiency of semantic judgments in terms of response times, consistent with distance-to-boundary models. Originally proposed to account for speed–accuracy tradeoffs in signal detection, these models predict that stimuli producing neural representations far from the classifier boundary yield quicker decisions (Pike, 1973; Ashby and Maddox, 1994). This phenomenon has also been demonstrated using neural distances measured with MEG and fMRI (Carlson et al., 2014; Ritchie et al., 2015; Grootswagers et al., 2018; Ritchie and de Beeck, 2019). Similarly here, more distinct semantic representations appear to facilitate rapid decision-making, suggesting that readable information in the brain may be used for behavior: the semantic system projects word meanings into separable activation patterns, which can be tapped into by downstream decision processes to speed responses. Notably, reaction times were longer for incongruent than congruent stimuli, consistent with prior psycholinguistic findings (Laver and Burke, 1993), indicating that contextual cues aid congruency judgments in both age groups. Thus, our results suggest that behavioral discriminability is driven by neural dissimilarity between semantic representations, not by conceptual semantic unrelatedness. In conclusion, decodability offers a framework linking behavioral performance to age-related neural alterations in semantic cognition.

Successful classification of age groups based on evoked contrast responses peaked between 300 and 600 ms, with ROC-AUC values reaching up to.8, indicating strong discriminative power of the logistic regression models during this time interval. This reflects the period of maximal divergence in neural activity patterns between age groups, highlighting a critical window for semantic processing where age-related changes are most pronounced, encompassing both the N400 and LPC timeframes. These findings align with a growing body of literature identifying mid-latency neural responses as particularly sensitive to cognitive aging (Kutas and Iragui, 1998; Federmeier and Kutas, 2005; Federmeier et al., 2010; Kemmotsu et al., 2012; Wlotko and Federmeier, 2012; Xu et al., 2017). This sensitivity likely reflects age-related neural reorganization and recruitment of alternative neural resources (Wlotko et al., 2010; Shafto and Tyler, 2014), potentially driven by the semanticization of cognition in older adulthood (Spreng and Turner, 2019).

## Conclusions

Our results suggest that the apparent inconsistencies across earlier psycholinguistic and imaging studies on aging and semantic cognition likely reflect methodological differences rather than true contradictions. Our findings help reconcile these perspectives by demonstrating that they can be concurrently valid: while the behavioral performance remains intact in older individuals, the spatiotemporal dynamics of neural processing can be altered. Specifically, the reduced N400 congruency effects in older adults aligned with electrophysiological evidence of weakened semantic priming, while increased frontal and bilateral activation mirrored fMRI findings of large-scale neural reorganization. Temporal decoding was able to capture differences in neural patterns between age groups, and cortical relevance patterns derived from the decoder weights further mirrored source modelling results. Decoding performance was directly correlated with reaction time, demonstrating a relationship between neuronal response patterns and behavior: increased neural separation between semantic concepts facilitated rapid decision-making. Overall, our findings suggest that healthy aging is associated with an adaptive reconfiguration of neural systems that maintains semantic proficiency despite broader age-related cognitive changes. Further investigation into the mechanisms of neural plasticity in late adulthood may facilitate the development of individualized approaches for sustaining linguistic and conceptual vitality throughout life.

## Acknowledgements

We acknowledge the following funding sources: Pietari Nurmi received funding from the Finnish Cultural Foundation, the Swedish Cultural Foundation in Finland, the Päivikki and Sakari Sohlberg Foundation, and the Finnish Brain Foundation. Heidi Ala-Salomäki received funding from the Jenny and Antti Wihuri foundation and the Finnish Cultural Foundation. Hanna Renvall received funding from the Academy of Finland (grant numbers 321460 and 355409 and the Flagship of Advanced Mathematics for Sensing Imaging and Modelling grant 359181). Mia Liljeström received funding from the Aalto Brain Center, the Finnish Cultural Foundation, the Päivikki and Sakari Sohlberg Foundation and the Swedish Cultural Foundation in Finland. This work was supported by the Helsinki-Uusimaa Regional Council. We are grateful to all the participants for taking part in the study.

